# *Arc* Regulates a Second-Guessing Cognitive Bias During Naturalistic Foraging Through Effects on Discrete Behavior Modules

**DOI:** 10.1101/2022.07.27.501779

**Authors:** Alicia Ravens, Cornelia N. Stacher-Hörndli, Jared Emery, Susan Steinwand, Jason D. Shepherd, Christopher Gregg

**Affiliations:** University of Utah, Department of Neurobiology, Salt Lake City, UT; University of Utah, Department of Human Genetics, Salt Lake City, UT; University of Utah, Department of Biochemistry School of Medicine, Salt Lake City, UT; University of Utah, Department of Ophthalmology & Visual Sciences, Salt Lake City, UT; Storyline Health Inc., Salt Lake City, UT

**Keywords:** foraging, learning, memory, *Arc*, behavioral economics, cognitive bias, decision heuristic, behavior cartography, PHATE, computational ethology

## Abstract

Foraging involves innate decision heuristics that are adapted for the wild but can cause economically suboptimal cognitive biases in some contexts. The mechanisms underlying cognitive biases are poorly understood but are likely genetic. Here, we investigate foraging in fasted mice using a naturalistic paradigm and uncover an innate “second-guessing” cognitive bias that involves repeatedly investigating an empty former food patch instead of consuming available food. Second-guessing prevents mice from maximizing feeding benefits in the task. Since learning and memory are involved, we tested roles for the synaptic plasticity gene, *Arc,* and found that *Arc^−/−^* mice show a specific lack of second-guessing. *Arc^−/−^* males reap benefits by increasing food consumption. Unsupervised machine learning decompositions of foraging show that *Arc* affects discrete, stereotyped foraging sequences that we call modules within a rich naturalistic behavioral landscape. Thus, our study reports a second- guessing cognitive bias, ethological roles for *Arc* in naturalistic foraging, and links between genetically determined foraging modules and cognitive bias in decision making.

## INTRODUCTION

Studies of animal foraging and human psychology have helped to uncover important rules and mechanisms of cognition and decision making, giving rise to the fields of behavioral economics and neuroeconomics ^1–5^. Economically optimal (or rational) behavior is operationally defined as behavior that maximizes the actor’s benefits. However, cognitive biases have been found that are systematic deviations from optimal economic decisions and involve innate heuristics that are “short-cut” rules the brain evolved for faster and more efficient decisions in the wild ^4–6^. Mechanistic studies of decision making typically focus on defining neurophysiological correlates, neuronal circuits, and mathematical models of decision processes ^7–17^. However, at least some cognitive biases have been observed in multiple different species ^1, 2,18^, suggesting evolutionarily conservation and a genetic basis. Indeed, given that cognitive biases are systematic, stereotyped behaviors proposed to involve innate fast- decision heuristics that are adapted for the wild, such biases likely point to components of behavior that are innate and genetically determined, rather than learned habits. So far, however, little is known about the genetic basis of the different cognitive biases affecting economic behaviors and foraging.

The study of genetically controlled behavioral components that evolved in the wild has been difficult due to the lack of precisely controlled paradigms that emulate natural conditions ^19^. Most lab studies focus on simplified and controlled binary choice tests that show limited richness in behavior ^20^. However, in nature, the *<u>entire behavioral sequence</u>* before, during, and after a choice shapes benefits and costs for natural selection. Approaches to study more naturalistic and rich behaviors have been lacking, though unbiased top-down decompositions of behavior are expected to improve our understanding of such complex behaviors ^21^. The development of data-driven methods to decompose spontaneous behaviors into discrete components is rapidly advancing this area of research ^22–24^. We recently developed a naturalistic foraging assay and unsupervised machine learning methods for top- down decompositions of foraging in mice to help reveal the structure and genetic mechanisms that affect different components of naturalistic foraging and decision processes ^25, 26^. Our paradigm mimics key elements of behavior in the wild, including a freely accessible home with sand patches of potential food, seeds, and digging. Using this approach, we found that foraging behavior involves finite, stereotyped and genetically controlled behavioral sequences that we call “modules” ^25^. Foraging modules are statistically reproducible from mouse to mouse and are identified from data describing decision and action sequences expressed during round trips from the home that can range from less than a second to hundreds of seconds in duration. With this approach, we are poised to investigate genetic mechanisms underlying different stereotyped components of naturalistic foraging.

Here, we deeply analyze foraging in fasted mice and uncover a cognitive bias that involves repeatedly investigating a former and now empty food patch rather than consuming readily available food. We refer to this as “second-guessing” because it involves repeatedly checking a former food patch after learning it doesn’t contain food. Second-guessing occurs in naïve male and female mice and contributes to suboptimal economic outcomes by failing to maximize food consumption. Given that memory confidence and repeated checks on memory accuracy have important roles in doubt and repetitive behaviors ^27–29^, we hypothesized that second-guessing is linked to memory and tested roles for the synaptic plasticity and memory gene, *Arc* ^30, 31^. Studies using standard lab tests of discrete behavioral responses, such as spatial memory ^33–35^, fear conditioning ^34, 36, 37^ and behavioral responses to novelty and stress ^38–40^ found that *Arc* is required for memory consolidation ^32^. However, ethological roles for *Arc* in rich, spontaneous naturalistic foraging have not been tested. We show that initial learning and memory responses are grossly intact in *Arc^−/−^* mice after 24 hours, but uncover a complete loss of second-guessing behavior that is replaced with increased food consumption in males. Using top-down, machine learning decompositions of the behavior, we uncover a finite set of stereotyped foraging modules that are affected by loss of *Arc* in a learned environmental context and underlie second-guessing versus feeding decisions. Thus, our study reveals a genetically determined “second-guessing” cognitive bias that is controlled by *Arc* affected foraging modules.

## RESULTS

### Identification of a “Second-Guessing” Cognitive Bias in Fasted, Foraging Mice

To investigate the nature of different memory responses during foraging, we designed a naturalistic paradigm in which mice have spontaneous access to a foraging arena and their home. Using methods from the foraging ecology field, our arena includes patches of sand, including one that has seeds. Mice learn the location of a food patch in the foraging arena during an Exploration phase (**Figure 1A**) and then 24hrs later, the mice are provided with access to the arena again during a Foraging phase, but the food is now in a different location (Pot4) (**Figure 1B**). Thus, we can test whether the mice express a memory response to the former food patch (Pot2) by measuring the time spent at Pot2 relative to patches that do not contain food (Pots 1 and 3). We also learn whether the mice discover and exploit the novel food patch (Pot4) by analyzing time spent at this location (**Figure 1B**), and ultimately, decompose the behavior using unsupervised machine learning methods to uncover foraging modules (see below). Overall, this paradigm enables us to investigate how memory of a former food patch and environment can shape foraging behaviors and decisions.

**FIGURE 1.**
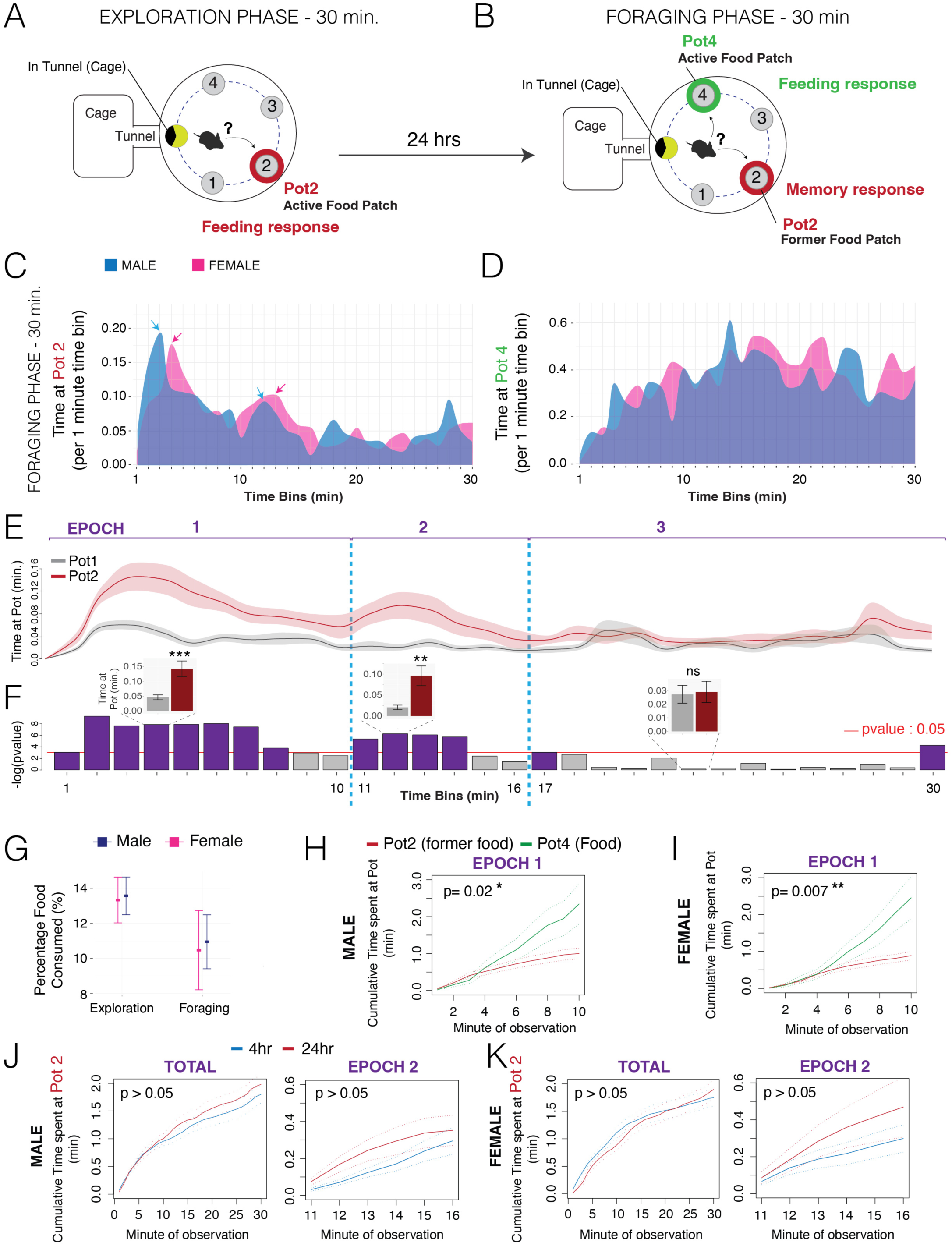
Mice show a discrete “second-guessing” behavioral epoch that involves repeatedly investigating a learned food depleted patch during naturalistic foraging. (**A** and **B**) Schematic summaries of the Exploration (A) and Foraging (B) phases of our naturalistic foraging assay. During the 30-minute Exploration phase (A), naïve mice learn Pot2 is a food-containing patch. During the 30-minute Foraging phase performed 24hrs later, the food patch has been moved and is now located in Pot4. The mice show a memory response to the former food location. (**C**) The plot shows the average time spent at the former food patch (Pot2) and the new food patch (Pot4) in the Foraging phase for male (blue) and female (red) mice (n=15). The time at each Pot is shown for 1 minute time bins over the 30-minute testing period. Male and female mice show an initial and then secondary peak of memory response behavior involving increased interactions with Pot2 (arrows point to peak interaction time points). (**D**) The plot below shows interactions with the new food patch (Pot4) and reveals a different pattern in which interactions increase steadily over the course of the testing period. (**E** and **F**) The top plots compare the mean time that mice spend at the former food patch (Pot2, orange) to time at a non-food containing control patch (Pot1, dark grey) during the 30-min Foraging phase (**F**, n=15; dark line shows the mean and shaded areas show the SEM). The bar plots in (**E**) show the p-value results for 1 minute sliding windows of a generalized linear model testing for a significant difference in time spent at Pot2 versus Pot1 in the male and female data (n=15). The purple bars show time windows that have a significant p-value after accounting for sex variance (p<0.05, Pot time ∼ sex + Pot type) and grey bars are windows that are not significant (p>0.05). The results reveal three discrete Epochs of memory response behavior (EPOCH 1, 2, and 3). Separation between epochs is defined as the transition from a non-significant effect (grey bars) to a significant Pot 2 bias (purple bars). The inserted bar plots show examples of the Time at Pot data for a 1-minute window in each epoch (grey bar is mean time at Pot1; red bar is mean time at Pot2; error bars show SEM). (**G**) The plots show the percentage of the total food that is consumed during the Exploration and Foraging phases by males (blue) and females (pink). The data show that the mice eat less than 20% of the total food, despite being fasted (n=15). Thus, the Epoch 2 memory response involving repeated searching of the former food patch (Pot2) in the Foraging phase is not caused by running out of food in Pot4. Mean ± SEM. (**H and I**) The plots show the cumulative time at the new (Pot4, green) versus former (P2, red) food patches for males (H) and females (I) during Epoch 1 of the Foraging phase. The data show that mice rapidly discover and value the new food patch during Epoch 1, showing significantly increased cumulative time at the new versus former patch (n=15, generalized linear model). Therefore, the Epoch 2 investigations of the former food patch (Pot2) are not caused by a failure to rapidly identify food. Mean±SEM. (**J and K**) The data show results of generalized linear model testing of whether the cumulative time spent at the former food patch (Pot2) over the entire Foraging phase (TOTAL) or during Epoch 2 only (EPOCH 2) is significantly affected by a 4hr versus 24hr memory consolidation gap between the Exploration and Foraging phases in male and/or female mice. Statistical results show that significant effects of the consolidation timing (timing) are not observed in total or in Epoch 2, nor are there significant interactions with sex. The plots show the data for males (J) and females (K). Mean±SEM, n=15.

We performed this foraging test on a cohort of wild-type adult male and female mice on a C57BL/6J background and first characterized interactions with the former food patch, Pot2, in the Foraging phase of the task. By analyzing the average time spent by all mice at Pot2 in time bins over the 30-minute testing period, we found that males (**Figure 1C, blue**) and females (**Figure 1C, Pot2, pink**) show 2 major peaks of interaction with the former food patch (**Figure 1C, blue and pink arrows**). The first peak shows the most Pot2 interactions over the trial, reaching a peak at ∼3 min in males and ∼4 min in females. This first peak of interaction then wanes, reaching a minimum interaction time at ∼10 min in males and 9 min in females (**Figure 1C**), which is then followed by a second peak at ∼12 min in males and 13 min in females (**Figure 1C, blue and pink arrows**). This second epoch of increased interaction wanes by 16 min in males and 17 min in females (**Figure 1C, blue and pink lollypops**). In contrast to interactions with Pot2, interactions with the new food patch, Pot4, show a different pattern and increase steadily, reaching a peak at ∼16 min that persists until the end of the assay (**Figure 1D**). Our results suggest that different epochs of interaction with the former food patch might exist that reflect discrete waves of memory-dependent behavioral responses and interactions with Pot2, and we therefore tested this.

To statistically test for discrete epochs of memory behavior, we performed a sliding window analysis of the data that involved sliding 1-min windows in which we compared time spent at the former food patch (Pot2) against time spent at a control patch (Pot1) that never contained food and is not close to the new food patch (Pot4) in the Foraging phase (**Figure 1E** and **F**). We analyzed the male and female data together using a model that absorbs the effect of sex and tests for a main effect of the Pot. From the results, we found 3 discrete epochs of memory-dependent response behavior in which the mice display waves of significantly increased time at Pot2 that are separated by windows where the mice do not spend significantly more time at Pot2 compared to Pot1 (Epoch 1: 0-10 min, Epoch 2: 11-16 min, Epoch 3: 17-30 min).

In Epoch 1, the animal’s increased interaction with the former food patch is rational given the initial expectation of food in this location. However, the Epoch 2 increase in interactions with Pot2 is an <u>economically suboptimal behavior</u> because the animal is fasted and knows that food is not located there but is abundant in Pot4. All mice have visited Pot4 by ∼2 minutes after entering the arena during the Foraging phase, with a mean latency of 25 seconds. We tested the possibility that mice perform this Epoch 2 behavior because they have consumed all the food in Pot4 and are now looking for more. However, we found that these fasted mice consume <20% of the total food available during the Exploration and Foraging phases of our task, which shows that they do not run out of food in Pot4 during the Foraging phase (**Figure 1G**). It is also not the case that the mice are failing to find the food in Pot4. We observed that they rapidly discover and show significantly increased interactions with the Pot4 food patch within the first minutes of the assay (**Figure 1H and I**). Collectively, the above results show that the Epoch 2 behavior is a discrete memory response during foraging that is directed at a former food patch that no longer contains food. Given the energetic costs versus caloric benefits for a fasted mouse to explore an empty food patch rather than consume available food, respectively, the Epoch 2 behavior shows evidence of being an innate, economically irrational cognitive bias to repeatedly “second-guess” a former food patch. We therefore refer to this behavior in Epoch 2 as “second-guessing”.

Second-guessing behavior could be affected by memory consolidation and the duration of the time separating the Exploration and Foraging phases of our assay. Here, we tested whether the duration of the consolidation period significantly affects second guessing by comparing a 4 hr (shorter-term memory) versus 24hr (longer-term memory) separation between the two phases. We found that the cumulative time at Pot2 over the total assay and during the Epoch 2 second-guessing behavior are not significantly different between the 4hr and 24hr conditions (**Figure 1J and K**). Therefore, second-guessing is not significantly affected by these shorter versus longer periods after training. We next tested if second-guessing is a genetically determined cognitive bias during foraging.

### *Arc* Knockout Mice Show Selective Loss of Second-Guessing with an Associated Increase in Food Consumption in Males

*Arc* regulates synaptic plasticity and long-term memory consolidation ^32^ and is a logical gene candidate for affecting second-guessing memory responses. Additionally, roles for *Arc* in naturalistic foraging have not been studied. Here, we profiled foraging in *Arc^−/−^* mice compared to ^+/+^ littermate controls in our assay. First, we tested how *Arc^−/−^* mice learn during the Exploration phase by examining how they adapt their behavior in the Foraging phase assay 24hrs later (**Figure 2A**). We found that <u>both</u> *Arc*^−/−^ and ^+/+^ mice show a significantly reduced latency to enter the foraging arena during the Foraging phase compared to the Exploration phase (**Figure 2B**). This shows that the ^−/−^ mice recall the general environment and adapt their behavior accordingly. We further found that both male and female *Arc^−/−^* and ^+/+^ mice show a significant preference for Pot2 within the first 5 minutes of the Foraging phase, indicating an initial memory to the former food patch (**Figure 2C and D**). Therefore, *Arc^−/−^* mice forage, learned the location of the food in the Exploration phase, and show initial memory-dependent behavioral responses 24hrs later in the Foraging phase.

**FIGURE 2.**
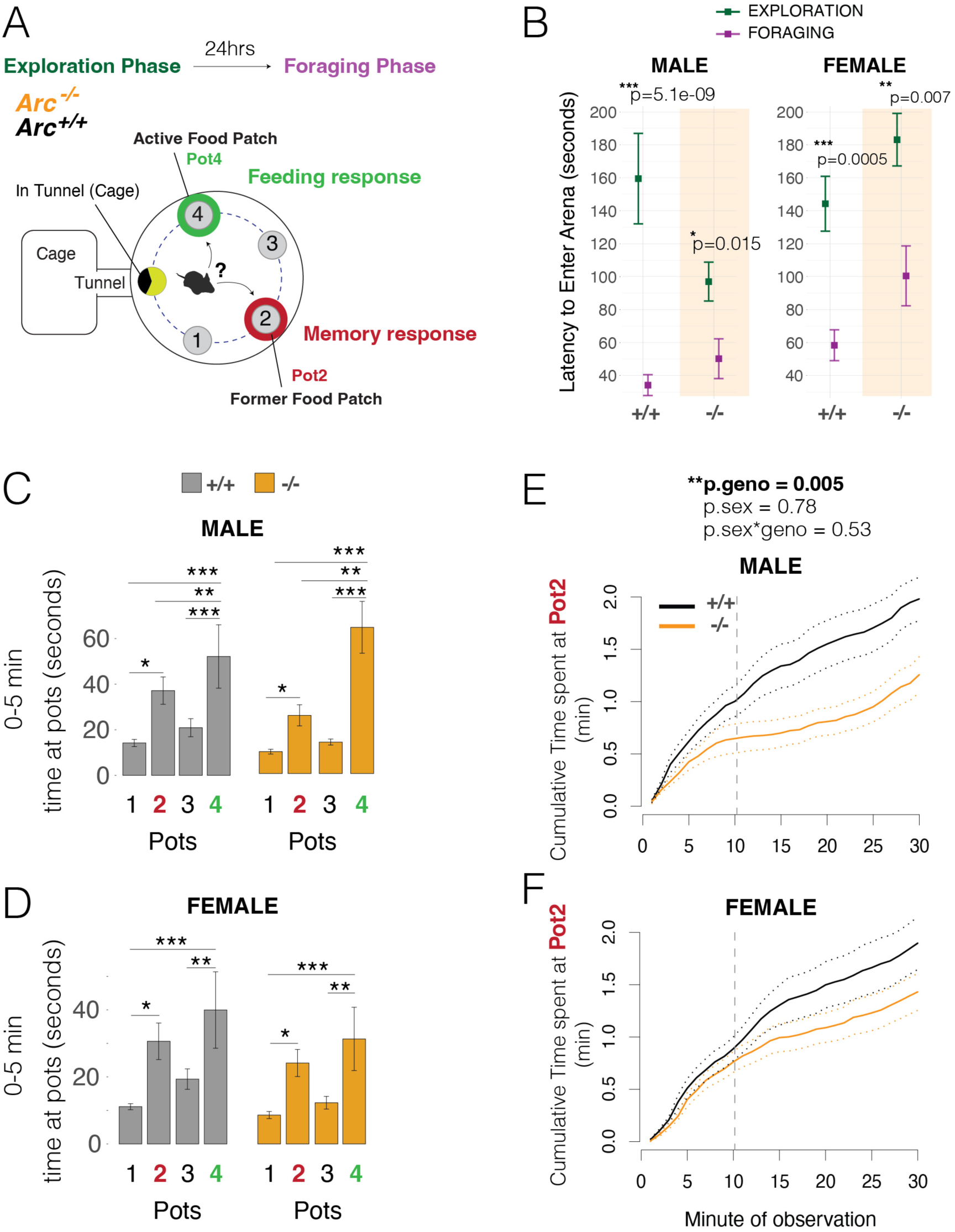
ARC knockout mice show grossly intact initial memory responses, but significant changes to memory responses during later epochs. (**A**) Schematic summary of experimental testing of naturalistic foraging phenotypes in *Arc^−/−^* versus *^+/+^* mice. The experiment tests behaviors related to exploitations of the available food-rich patch (Pot4) versus now food-absent former food patch (Pot2). (**B**) The plots show the latency to enter the foraging arena from the home during the Exploration (blue) versus Foraging (purple) phases for *Arc^−/−^* versus *^+/+^* littermates. Male and female *Arc^−/−^* and *^+/+^* mice show a significantly reduced latency to enter the arena in the Foraging phase compared to the Exploration phase (n=15). t-test. Mean ± SEM (**C** and **D**) The bar plots show the total time spent at each of the pots during the first 5 minutes of the Foraging phase for male (C) and female (D) *Arc^−/−^* mice (orange) and *^+/+^* (black) littermates. *Arc^−/−^* and *^+/+^* littermates spend significantly increased time at the former food patch (Pot2), showing they learned and recall this former food patch, and at the new food rich patch (Pot4). A significant main effect of Pot was observed (p<0.0001). The +/+ and −/− mice are not significantly different (p>0.05). N=15, Mean ± SEM, one-way ANOVA with Tukey post-test. *p<0.05, **p<0.01, ***p<0.0001 (**E** and **F**) Plots of the cumulative time at Pot2 during the Foraging phase show that *Arc^−/−^* males (E) and females (F) exhibit later epoch behavioral changes compared to +/+ controls by spending less time at Pot2 beginning at ∼10 min into the Foraging phase (grey dashed line). P-values from generalized linear modeling of cumulative time at 30 min are shown and indicate a significant main effect of genotype (p.geno), but no effect of sex (p.sex) or interaction (p.sex*geno). N=15, Mean ± SEM.

By analyzing plots of the cumulative time spent at Pot2 over course of the assay, we found a significant genotype effect in which ^+/+^ males and females continue to accumulate increased time at Pot2, but ^−/−^ mice exhibit a significant reduction in Pot2 interactions that becomes pronounced after approximately 8 minutes into the assay in males (**Figure 2E**) and 15 minutes in females (**Figure 2F**). This finding shows that *Arc* most strongly affects the behavior during later epochs, rather than the initial memory responses to the former food patch, Pot2.

The memory phenotype in *Arc^−/−^* mice could be secondary to changes in motivation to feed. To test this, we examined food consumption during the Exploration phase, which lacks a memory component and found that *Arc^−/−^* mice do not consume less food or spend less time at the food (**Figure S1A and B**). This indicates that their drive to feed is not significantly reduced compared to ^+/+^ mice in the assay. Alternatively, *Arc^−/−^* mice might be slow to discover the novel food patch in the Foraging phase and therefore express a delayed, but elevated interest in Pot4 over Pot2. However, we found that *Arc^−/−^* mice discover and significantly increase Pot4 interactions within the first ∼8 minutes of the Foraging phase like ^+/+^ mice (**Figure S1C**). Given that *Arc^−/−^* mice do not display overt reductions to the drive to consume or find food, we concluded that they display bona fide changes to later memory response epochs during foraging, rather than changed motivations to feed. We therefore directly tested second-guessing behavior.

To examine roles for *Arc* in second-guessing, we visualized the pattern of interactions with Pot2 over the course of the 30 min Foraging phase in 1 minute time bins to reveal memory-dependent epochs (**Figure 3A and B**). We found that *Arc^−/−^* males show a specific loss of the Epoch 2 second guessing behavior, as they no longer show a second wave of interest in Pot2 (**Figure 3A**). Similar, albeit less robust effects were observed in females (**Figure 3B**). We further tested this difference by computing the cumulative time at Pot2 during Epochs 1, 2 and 3 in the *Arc^−/−^* versus *^+/+^* mice. We found a significant main effect of genotype (^−/−^ versus ^+/+^) in which *Arc^−/−^* mice show a loss of Pot2 interactions during Epoch 2, but not Epochs 1 or 3 (**Figure 3A, B;** see statistical results in top boxes). Thus, *Arc^−/−^* mice show selective loss of second-guessing during foraging.

**FIGURE 3.**
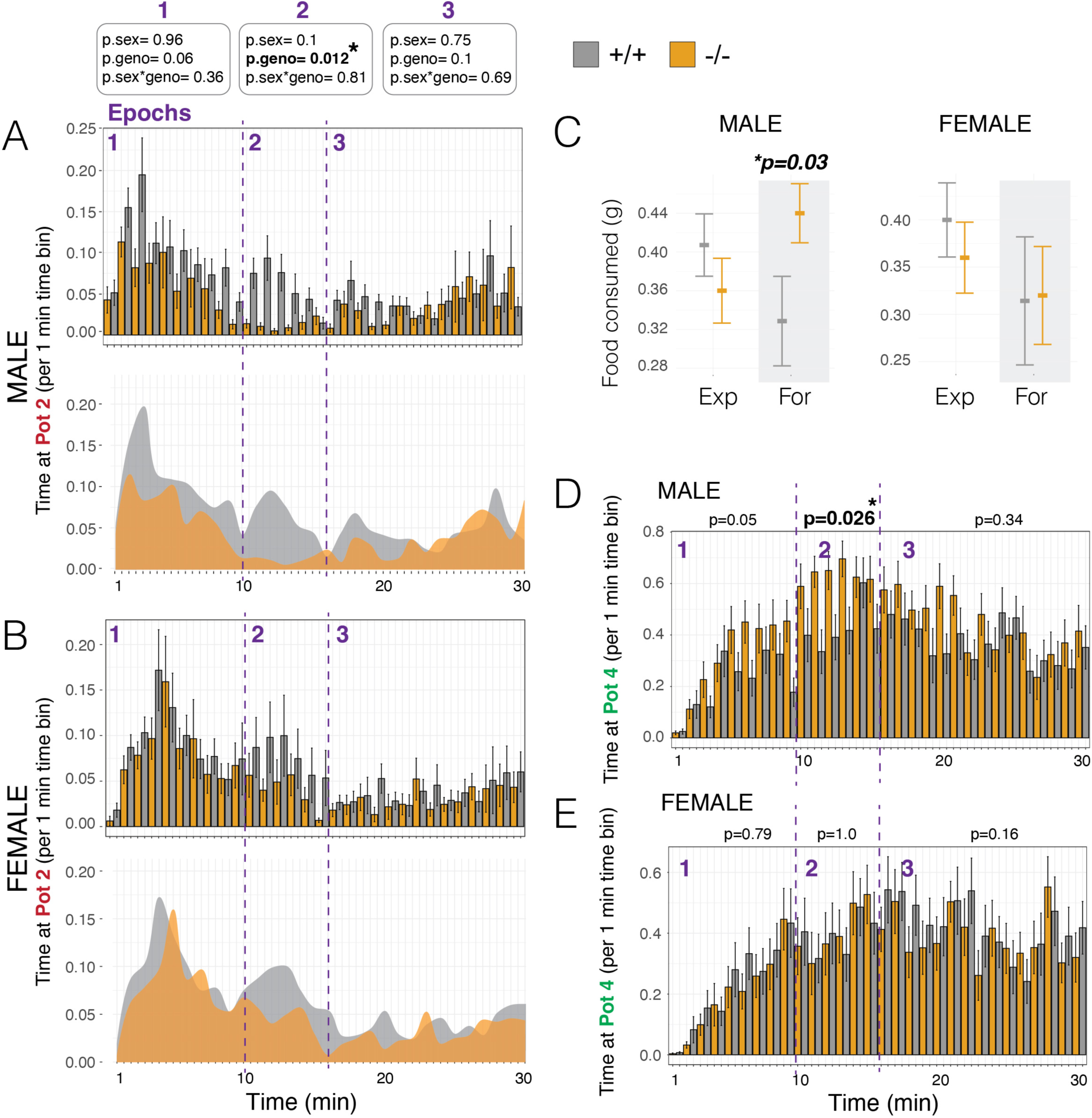
*Arc^−/−^* mice show deficits in second-guessing behavior. (**A** and **B**) The barplots show data for the time spent at the former food patch (Pot2) in the Foraging phase in 1 minute time bins for *Arc^−/−^* (orange) males (**A**) and females (**B**) compared to +/+ littermate controls (grey). The 1, 2 and 3 memory response epochs are delineated by purple numbers and dashed lines. The data show significant loss of the epoch 2 second-guessing memory response in *Arc^−/−^* males and females. Significant changes to Epoch 1 and 3 behaviors were not observed. Plots below summarize the phenotypic effect by showing the mean time for knockouts versus wildtypes at Pot2 over the Foraging phase (grey). P-values for generalized linear model are shown on the top. (n=15) (**C**) The plots show the total food consumed by male and female mice in the Exploration (Exp) phase and Foraging (For) phase for *Arc^−/−^* (orange) mice compared to +/+ littermate controls (grey). Males show significantly increased food consumption in the Foraging phase. N=15, Generalized linear model interaction for phase and food consumed. (**D** and **E**) The barplots show data for the time spent at the new food patch (Pot4) in the Foraging phase in 1 minute time bins for *Arc^−/−^* (orange) males (**F**) and females (**G**) compared to +/+ littermate controls (grey). Male *Arc^−/−^* mice show increased time at the food patch specifically during the Epoch 2 second-guessing memory response. N=15, Mean ± SEM.

Given the absence of the second-guessing response, we determined what behaviors the *Arc^−/−^* mice are doing in its place. We found that male KO mice show significantly increased food consumption in the Foraging phase (**Figure 3C**) and shift to significantly increased interactions with the Pot 4 food patch during the Epoch 2 memory response (**Figure 3D**). This effect was not observed in females (**Figure 3C and E**) and deeper analyses of the female data found that they did not spend significantly increased time at Pots 1 or 3 or in the home. Thus, *Arc^−/−^* females appear to redistribute their behavioral activity broadly during Epoch 2 rather than focusing on a specific patch or location in the arena. Overall, we uncovered a role for *Arc* in affecting second-guessing during foraging and in the absence of second-guessing *Arc^−/−^* males show more economically optimal behavior by increasing food consumption.

### *Arc* Affects a Discrete Set of Foraging Modules that Determine Second-Guessing

Previously, we found that naturalistic foraging behavior in mice shows evidence for having a modular structure and involve finite, stereotyped and genetically determined behavioral sequences ^25^. Here, we tested whether second-guessing involves modules of foraging behavior that are affected by *Arc*. To first uncover the modules expressed by the *Arc^−/−^* and *^+/+^* mice, we applied our DeepFeats algorithm to decompose, describe and classify the behaviors expressed by the mice (n=60) ^25, 41^. For each of the 1,609 round trip foraging excursions expressed by the mice, we captured 57 diverse measures of how the mice move and where they go while foraging. Principal component analysis (PCA) was applied for data dimension reduction in combination with the DeepFeats behavioral module detection algorithm. The data was partitioned into balanced training and test datasets of mice and our analysis found that 8 principal components (PCs) best reveal candidate modules in the training data (**Figure S2A,B**). With these PCs, we found 43 clusters of different foraging sequences that reflect candidate foraging modules (**Figure S2C**). With a modified in-group proportion permutation test, we then evaluated the reproducibility of the training dataset clusters in the test dataset (**Figure S2C,D**) and found that 24 of the 43 clusters are significantly reproduced in the test set, which we defined as significant, stereotyped modules (**Figure S2D**). Overall, 942 of the 1,609 total round trip foraging excursions are classified in modules, while the remaining excursions are non-modular and reflect more stochastic and idiosyncratic behaviors.

Next, we tested whether loss of *Arc* impacts the expression of specific modules, or alternatively affects non-modular behaviors. A generalized linear model testing for an interaction effect between *Arc* genotype and the expression frequencies of different modules (24 different modules) and non-modular behaviors (19 non-modular excursion clusters) revealed that the most significant genotype X behavior type interaction occurred for modules in the Foraging phase (p=2.7×10^−11^; **Figure 4A**, dark bars). Post-tests with q-value corrected p-values revealed that the expression frequencies of Modules #1, 2, 10, 12, 21 and 43 are significantly affected in *Arc^−/−^* mice (q<0.05), while the other 18 modules are not significantly affected. In our examination of non-modular behavior types, we did not observe a significant interaction in the Foraging phase and found a modest effect in the Exploration phase (p=0.04; **Figure 4A**, light bars). We further confirmed that *Arc* predominantly affects module expression in the Foraging phase with a different analysis that tested for changes to the stereotyped expression order of modules by the mice, which we found to be significantly changed in *Arc^−/−^* mice in the Foraging phase, but not the Exploration phase (**Figure S3A-F**). In conclusion, loss of *Arc* most strongly affects the expression of modules, rather than non-modular excursions, and impacts a specific subset of modules in the Foraging Phase (**Figure 4A**).

**FIGURE 4.**
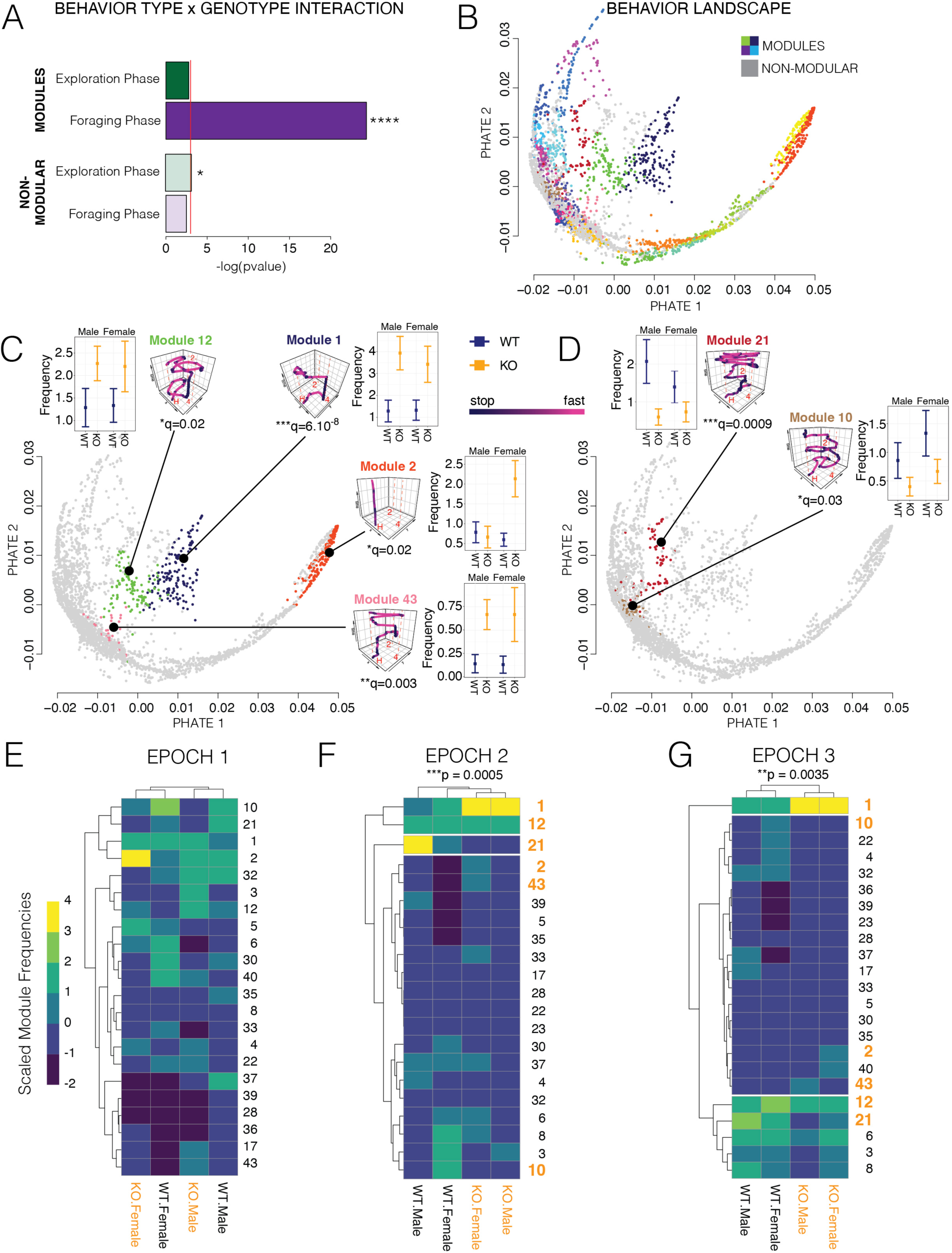
Loss of *Arc* affects the expression of discrete foraging modules that underlie second-guessing versus feeding behaviors. (**A**) The dark bar plot shows the p-values for a generalized linear model testing the interaction effect between genotype and module on module expression frequency data for male and female *Arc^−/−^* and ^+/+^ mice in the Exploration and Foraging phases (n=15). The light bar plot shows the results for excursion behavior clusters that are not statistically significant modules where the interaction effect is tested for genotype and behavior cluster. The results show that Foraging phase modules are the most strongly affected and have a significant genotype X module interaction effect indicating that a subset of modules are significantly impacted by loss of *Arc.* *p<0.05, ****p=2.7×10^−11^ (**B**) The plot shows a 2D PHATE projection of the multi-dimensional data for each excursion expressed by the male and female *Arc^−/−^* and ^+/+^ mice in the study (1609 excursions shown). The excursions classified as significant foraging modules are colored and non-modular excursions are grey. The results show the structure and relationships of the different behavioral components of naturalistic foraging (see Supplemental Video S1 and S2 for full 3D projection). (**C** and **D**) The plots show the 2D PHATE projection of the excursion data where the colored excursions indicate modules with significantly increased (C, Modules 12, 1, 2 and 43) versus decreased (D, Modules 21 and 10) expression in *Arc ^+/+^* and *^−/−^* mice. Representative X-Y coordinate traces of the movements over time (Z-axis) are shown for each significantly affected behavior module, which show the center point tracking data for excursion closest to the centroid of the cluster (see **Figure S4** for more details). Traces are colored by movement velocity (Black is slow or stopped and pink is moving). Red lines and letters indicate the home tunnel (H), Pot2 former food location (2), and Pot4 current food location (4). The plot inserts show the expression frequency data for each affected module for male and female *Arc ^+/+^* (WT, blue) and *^−/−^* (KO, orange) mice. A linear model testing for a main effect of genotype was applied to the data for each module and multiple testing was controlled using the q-value (shown below trace). The excursion data in the PHATE map is highlighted for each *Arc* affected module to show grouping structure and distance relationships in space (see main text). Mean ± SEM. *q<0.05, **q<0.01, ***q<0.001. (**E-G**) The heatmaps show the aggregated expression frequency of different foraging modules (rows) in male and female *Arc* KO (−/−) and WT (+/+) mice (columns). The data has been scaled by column (see legend). Unsupervised hierarchical clustering of the data grouped male and female KOs separately from WTs in memory response Epochs 2 (E) and 3 (F), but not Epoch 1 (G). A chi-square test on the Epoch 2 and 3 datasets showed statistical significance (p-value shown on top), indicating significant dependence of genotype on module expression. Modules that are significantly affected in *Arc−/−* mice are highlighted in orange numbers, while modules that are not significant are in black type. The data reveal specific modules that are most strongly affected, including Module 1, which has increased expression in *Arc−/−* mice during Epochs 2 and 3.

To visualize the global structure of naturalistic foraging in mice and learn the relationships between different modules and non-modular behaviors, and between *Arc* affected and unaffected modules, we visualized our entire multi-dimensional behavior dataset for all 1609 foraging excursions in our data using PHATE (Potential of Heat Diffusion for Affinity-based Transition Embedding) ^42^. PHATE is an unsupervised algorithm for 2D and 3D embedding of high dimensional data that uncovers accurate, denoised local and global data structures ^42^. For our study, the results show the global landscape of foraging behavior in which each point is an embedding of the data describing a single round trip excursion from the home (**Figure 4B**). By annotating excursions in the 24 modules (**Figure 4B, each module has a different color**) and the remaining non-modular excursions (**Figure 4B, grey dots**) in this map, we found that the excursions for different modules are well separated, and individual excursions assigned to the same module are grouped, rather than intermingled, as expected. This independent approach therefore confirms the classification of these discrete behaviors. The map also reveals the relationships between different modules since modules that are more similar are closer and connected to each other in space. The branches and connections between modules in the diffusion map show the global structure and continuum of mouse foraging. Supplemental videos of rotating 3D PHATE projections of modules and non-modular excursions (**Supplemental Video S1**), and modules alone (**Supplemental Video S2**), show the structure more clearly.

We visualized the data for the subset of modules that are significantly affected by loss of *Arc* (**Figure 4C,D** and **Supplemental Video S3**). The PHATE map shows that these affected modules involve well separated and distinct clusters of excursions (**Supplemental Video S3**) and a heatmap shows how each retained principal component differentiates the modules (**Figure S4A**). Modules 1, 2, 12 and 43 have significantly increased expression in *Arc^−/−^* mice (**Figure 4C,** graph insets), while Modules 10 and 21 show decreased expression (**Figure 4D,** graph insets). Module 2 is distinctive, involving a short dart into and out of the foraging arena that is significantly increased in *Arc^−/−^* females (**Figure 4C, orange**). Modules 1, 12 and 43 involve reduced exploration and more extended visits to the food patch (Pot4) (**Figure 4A**, insets and **Figure S4B,C).** Module 1 is the most significantly increased in *Arc^−/−^* mice and is a simple decision and action sequence that involves going directly from the home to the food patch (Pot4) and then returning to the home (**Figure 4C**, Module 1 trace inset, and **Figure S4B,C**). Modules 10 and 21 show decreased expression by *Arc^−/−^* mice and representative traces of the decision and action sequences for these modules reveal that they are complex exploratory behaviors that involve increased interactions with the former food patch (Pot2), as well as other components of the environment (**Figure 4D**, Module traces and graph insets and **see Figure S4D,E**). Our results show that *Arc* affects the landscape of naturalistic foraging behavior by impacting discrete modules. *Arc* likely influences second-guessing by affecting the expression of these modules in particular epochs of the behavior, which we investigated next.

To begin, we tallied the expression frequency of each module for the *Arc^−/−^* and ^+/+^ males and females for Epochs 1, 2 and 3 of the Foraging phase. The results are presented in heatmaps scaled by column, which show modules with relatively high versus low expression frequencies for each sex and genotype (**Figure 4E-G**). Unsupervised hierarchical clustering analysis of the module expression data found differences between the *Arc* ^−/−^ and ^+/+^ mice in Epochs 2 and 3 with statistically significant row by column dependency (**Figure 4E,F;** Fisher’s Exact Test). In contrast, this effect was not observed in Epoch 1 where the columns cluster by sex instead of genotype (**Figure 4G**). The preference for *Arc^−/−^* mice to express Module 1 over Modules 10 and 21 becomes pronounced during Epoch 2 and persists during Epoch 3 (**Figure 4F and G**), which supports the conclusion that changes to the expression of these discrete modules during Epoch 2 contributes to the switch from second-guessing (Modules 10 and 21) to increased food consumption (Module 1) in *Arc^−/−^* mice. Overall, we mapped the effects of *Arc* in naturalistic foraging to a second-guessing cognitive bias involving specific decision and action sequences expressed during a defined temporal epoch of the behavior.

## DISCUSSION

Cognitive biases are systematic behavioral biases that can result in decisions that are not economically optimal in some contexts. Their mechanistic basis is poorly understood, but the conservation of some cognitive biases across different species suggests they may be genetically determined. Here, we uncovered a second-guessing cognitive bias in fasted foraging mice that involves repeatedly visiting an empty former food patch instead of consuming available food to maximize caloric benefits. This bias occurs in both sexes and shows significant reproducibility across different mice, indicating that second-guessing is an innate behavioral bias rather than a learned habit. The bias is strongest in males. *Arc^−/−^* mice show specific loss of second-guessing, and in its absence, *Arc^−/−^* males reap the benefits of more economically optimal behavior by increasing food consumption. *Arc* affects second-guessing by impacting the expression of specific foraging modules within the landscape of behaviors that comprised naturalistic foraging in our assay. The affected modules are discrete, stereotyped decision and action sequences that shape interactions with the available food patch versus second-guessing of the former food patch. While *Arc^−/−^* mice lack second-guessing, their initial memory responses that reflect learning and recall of the environment are grossly intact. Thus, our study reports a genetically determined cognitive bias from studies of naturalistic foraging and an ethological role for *Arc* in regulating this second-guessing bias through effects on the expression of a subset of foraging modules during a specific epoch of the behavior. Our findings provide a rationale for future efforts aimed at defining links between genetic mechanisms, naturalistic behavior modules and cognitive biases.

Cognitive biases are thought to be related to innate decision heuristics that evolved for efficient decision processes in an organism’s native environment ^43^. While such biases can be economically suboptimal in some contexts and contribute to poor decision making, they likely evolved to be adaptive and advantageous in the native context ^6, 44^. It has been pointed out by others that behavioral ecologists often ignore the potential for genetic factors to constrain animal decision making ^45^. This is unfortunate since decision biases and constraints likely point to important roles for genetic factors that evolved to shape natural behavior. In support, we show that second-guessing is cognitive bias that is expressed by mice during naturalistic foraging and that this bias has a genetic basis. *Arc* is required for second-guessing and since *Arc^−/−^* males show increased food consumption in place of second-guessing, our data reveal that this cognitive bias can be genetically modified to yield more economically optimal behavior in our task. Second-guessing likely has important adaptive advantages in the wild. If food patches can appear, disappear, and reappear quickly, then it may be important to have constantly updated knowledge of different food patches in the local environment and the dynamics of food availability, and selection would lead to the development of discrete behavior patterns that solve this problem. Moreover, if animals must frequently abandon available food due to high risks of predation or encounters with conspecific competitors, then updated knowledge of alternative patches and frequent patch switching could be important for survival. Thus, second-guessing may be statistically and economically rational in the wild even when a known food patch is available.

While both males and females display *Arc*-controlled second-guessing, we observed sex differences that could involve sex hormones, metabolism, or other mechanisms yet to be determined. The second-guessing bias is strongest in males and while *Arc^−/−^* male mice consume more food in the absence of second-guessing, this effect is not significant in females. Additionally, *Arc^−/−^* females uniquely show significantly increased expression of foraging Module 2, which involves brief darting into and out of the foraging arena from the home. Thus, we uncovered multiple sex differences in how *Arc* affects particular foraging decisions in males and females. While sex differences in decision-making heuristics are debated ^48, 49^, it is known that risk taking, impulsivity and behavioral prioritization differ between males and females ^48^, which could contribute to sex differences in second-guessing. Indeed, our other foraging studies also found important sex differences in mice ^25, 26^.

We provide evidence that *Arc* impacts second-guessing and foraging behavior by affecting the expression of discrete foraging modules, rather than more non-modular and free behaviors. Both *Arc^−/−^* males and females show increased expression of Module 1 and decreased expression of Modules 10 and 21, revealing shared behavioral effects at the module level across the sexes. Module 1 involves a direct excursion from the home to the food patch and back, while Modules 10 and 21 involve more repeated visits to the former food patch (Pot2) and complex exploratory behavior. We visualized the global structure of mouse foraging in our assay, which revealed discrete modules affected by *Arc* within this map. Our results show the potential for unsupervised computational approaches to “behavioral cartography” that illuminate maps of complex naturalistic behavior and how different genetic factors affect specific components. *Arc* affects the expression of discrete foraging modules that shape second-guessing versus feeding during later epochs of foraging. Our sliding window analysis showed that these epochs are discrete temporal phases of memory-response behavior and transitions between each epoch occur when time spent at the former food patch (Pot2) drop is not significantly different from time spent at a control patch that never previously contained food (Pot1).

*Arc* is often considered a general regulator of synaptic plasticity and memory, and standard lab memory tests indicate that *Arc^−/−^* mice fail to consolidate long-term memories ^34, 36, 50^. Our foraging results show that initial Epoch 1 memory responses are grossly intact in *Arc^−/−^* mice, while Epoch 2 second-guessing and later memory epochs are more strongly affected, providing insights into ethological roles for *Arc* during foraging. The second-guessing effects of *Arc* could be due to changes in cognitive flexibility and perseveration, which have been reported in *Arc* transgenic mice where disruption of Arc protein degradation led to abnormal synaptic plasticity and behavior ^35^. *Arc^−/−^* mice may also have less robust memory that affects second-guessing during Epoch 2 and 3 but has little gross effect during Epoch 1. Overall, our foraging study indicates that *Arc*’s role in learning and memory are more task, context, and sex-specific than has been previously observed. Future studies are needed to define the brain regions, neural circuits, and types of cellular plasticity mechanisms in which *Arc* regulates second-guessing in males and females using conditional genetic approaches.

We previously showed that mouse foraging involves finite, stereotyped behavior modules that are segmented by round trip excursions from the home ^25, 26^. Here, using PHATE projections of multi- dimensional foraging data describing individual excursions, we showed that these modules are well defined components of the naturalistic foraging behavior landscape and are differentiated from more stochastic and free non-modular behaviors. Foraging modules share some conceptual similarities to other stereotyped movements, such as motifs or syllables ^22, 51^, but are more complex, compound decision and action sequences that incorporate both movement and contextual information. Here, we showed a link between specific foraging modules and an economically irrational cognitive bias, and that loss of *Arc* selectively eliminates the cognitive bias. Over 50 different cognitive biases have been described in humans ^46, 47^. Based on our results, we speculate that foraging modules have important links to different cognitive biases and innate decision heuristics. Indeed, modularity is central to phenotype evolution by enhancing evolvability, such that existing modules can be reorganized, duplicated, and modified without recreating their constituent parts and structure *de novo* ^52^. While modularity is a component of all biological systems ^52–55^, it is a newer concept in behavioral economics. Stereotyped, hierarchical and modular behavior has been described in *C. Elegans, Drosophila* and mice ^24, 25, 56–62^. Our results show that *Arc* affects the expression frequency of specific foraging modules in a context and time dependent manner, and we expect that other genetic mechanisms yet to be discovered affect the form, timing, and expression frequency of second-guessing modules. Second- guessing could share mechanisms with doubt and obsessive-compulsive behaviors, making further dissections of the mechanisms involved of potential biomedical importance ^29, 63^.

## LIMITATIONS OF THE STUDY

Our study used fasted mice to study the cognitive bias of investigating former food patches versus eating available food. We have not determined how different internal metabolic states affect second- guessing, which limits our understanding of when and how this bias arises. Moreover, our focus on fasted mice differs from published studies of *Arc^−/−^* mice that reported loss of long-term memory consolidation and may contribute to differences in which we observe that some initial memory responses are intact in *Arc^−/−^* mice. It is also possible that memory consolidation in this naturalistic foraging task may take longer than 24hrs and thus may only be Arc sensitive over longer periods between exploration and foraging phases that we didn’t test.

## ACKNOWLEDGEMENTS

We thank members of the Gregg and Shepherd labs for their critical reading of the manuscript and input on the study. This work was supported by funding from the National Institutes of Health (R01AG064013, R21MH120468, R01MH109577, and R21MH118570 grants to C.G., and R01MH112766 to J.D.S.).

## AUTHOR CONTRIBUTIONS

A.R. performed the behavioral experiments in the *Arc* knockout mice. C.S.H. performed the data analysis and made the figures. J.E. and C.G. developed the behavior analysis software and algorithms. C.G. and J.D.S. wrote the manuscript with input from all authors, supervised the study design, methods and data interpretation and provided funding.

## METHODS

### RESOURCE AVAILABILITY

#### Lead Contact

- Further information and requests for resources and reagents should be directed to and will be fulfilled by the Lead Contact, Christopher Gregg.

#### Materials Availability

- Datasets generated in this study are available from the Gregg lab.

#### Data and Code Availability

- All datasets and original code are available from the lead contact upon request.
- Any additional information required to reanalyze the data reported in this paper is available from the lead contact upon request.

### EXPERIMENTAL MODEL AND SUBJECT DETAILS

#### Mice

##### Housing and husbandry

All experiments were conducted in compliance with protocols approved by the University of Utah institutional animal care and use committee (IACUC). C57BL6/J and *Arc/Arg3.1* knockout mice were bred and housed on static racks in the Biopolymers Building near the lab’s behavioral testing room on a 12hr reversed light, 11pm on and 11am off. The generation of the *Arc/Arg3.1* knockout mouse line has been previously described ^34^. All mice were given water and food (Harlan-Teklad 2920X soy protein-free) *ad libitum*, with the exception of a single overnight fast for mice tested for foraging behavior (see METHOD DETAILS, *Behavior*, foraging). Adult breeders (6 weeks to 1 year of age) were paired continuously, and pups were weaned at postnatal day 21 (P21) and cohoused with up to five same-sex littermates or similar aged same-sex mice of the same line; mice were never singly housed. Ear punches were taken at P21 for both genotyping biopsy samples and mouse identification purposes. Genotyping was performed by Transnetyx.

##### Genotyping

Ear punches taken at P7-P21 were lysed in 75µL of 25mM NaOH + 0.25mM EDTA with a 1 hour incubation in a thermalcycler at 98°C. Lysates were then pH neutralized with an equal volume of 40mM Tris.HCl, pH5.5. Two µL of lysates were then added to make 20µL PCR reactions with DreamTaq Green Master Mix (ThermoFisher, Cat# K1081) and 0.5µM primers.

### METHOD DETAILS

#### Foraging

Foraging behavior testing was performed as detailed previously ^25^. In brief, in preparation for the foraging assay, mice were first habituated with sand (Jurassic play sand, Jurassic Sand) and seeds (Whole millet, Living Whole Foods) for two days in their home cage. On day one, seeds are spread on top of sand in the bottom of a Petri dish and the dish is placed on the bedding in the home cage for the mice and pups to explore. On day two, seeds are covered with sand in the bottom of the Petri dish in the home cage for the mice and pups to dig in and explore. To motivate animals to feed, mice were food deprived prior to testing to achieve 8%–10% weight loss at the time of testing. We selected this weight loss target after several pilot studies with the goal of achieving some consistency in the motivational states of the animals at different ages and not compromising health or activity. To achieve the intended weight loss and motivational state, adult mice were food deprived for 24 hours. Water is available *ad libitum* at all times except when mice are in testing cage (2 × 1 hour).

Mice are housed in a room with an 11:00 – 23:00 dark cycle, so that testing is performed during the dark cycle. For testing, mice are moved into the behavior room prior to the start of testing for at least 1 hour for habituation to the new room. All testing is performed in the dark and video recording is done using infrared illumination and all manual procedures are done in the dark using red light. The mouse to be tested, and their home cage soiled bedding, are moved to the testing-cage and allowed to habituate. At the start of testing, the testing-cage is attached to the arena via the tunnel, the mouse now has access to the arena and video recording starts for the Exploration phase. Mouse behavior is recorded continuously during the 30 min Exploration phase trial under infrared lights. Noldus Ethovision software v14 were used for video tracking. After completing the Exploration phase, the mouse is returned to the testing cage with water but no food until the Foraging phase 24 hours later. For the Foraging phase, the testing cage is then gently attached to the tunnel and access to the arena is possible and the Foraging trial begins. Video recording of the Foraging phase is performed for 30 minutes. After testing, mice are placed in a new cage with food and are returned to the mouse colony room. Between each Exploration and Foraging phase trial, the entire arena, including walls, platform, tunnel and steel pots, are wiped clean with 70% ethanol.

##### Preparation of Sand and Seed Pots for the Foraging Assay

Three stainless steel pots (Resco, diameter 5.5cm, depth 4cm) were filled with 95 g of sand. For the Exploration phase, 1 pot is filled with 80 g of sand covered with 2.5g of seeds. On top of seeds, a layer of 12 g of sand is added to cover seeds. This sand is then covered with 0.5g of seeds. This pot is placed in position 2 in the arena. For the Foraging phase, 1 pot is filled with 80 g of sand, 3g of seeds on top of sand and additional 12 g of sand to cover all seeds. This pot is placed in position 4 in the arena. All pots are weighed before and after the trial to measure the sand displaced from each pot. Remaining seeds and hulls left in the pot and on the platform are measured after each Exploration or Foraging trial to determine the amount of seeds consumed by the mouse during the trial. Used sand is collected after every trial and set aside. At the completion of all testing, the used sand is autoclaved before reuse in future trials.

##### Foraging Arena Construction

The foraging arena, tunnel and testing cage were custom built with acrylic plastic (Delvies Plastics, Salt Lake City, UT, USA). The 14 cm long tunnel enters the platform from underneath through one of the five 5.5 cm diameter holes in the arena platform. The platform is 8 cm above epoch level and is made from 0.5 cm thick white Plexiglas. The arena is made from a transparent Plexiglas tube and the 0.4 cm thick walls raise 42 cm above the platform. The arena has a diameter of 35 cm. The walls of the arena were roughened with sandpaper to limit glaring and recording artifacts.

##### Arena Tracking Zones

The arena is organized into zones that are used to breakdown the behavior and foraging strategies used by each animal in the assay. The arena is divided into five sectors and the boundary of each sector is the midpoint between two pots. The arena is further divided into three concentric circles, including the middle center zone, the intermediate zone and the outer wall zone. The outer radius of the *Intermediate* zone intersects with the center of the pots and tunnel entry. The radius of the *Center* zone is half the radius of the *Intermediate* zone. A zone is also created around each pot in the arena. *Pot* zones have a radius of 1.7x the radius of the pot itself. Finally, to learn about the behavior of the animal related to entries to and exits from the arena, we define zones around the tunnel entry. The *Tunnel Entry* zone aligns with the entry hole of the actual tunnel. The *In Tunnel* zone is covering the most peripheral area of the *Tunnel Entry* and tracks the mouse just before leaving the arena completely. Whenever the mouse is in the cage, the tracking system is recording the mouse as being in the *In Tunnel* zone. The *Tunnel Zone* area has the same radius as the *Pot* zones.

##### Automated Tracking

At the start of the trial, the tracking begins with a 10 s delay to allow time for the connection of the testing-cage to the arena. The mouse is first tracked when it appears in the *In Tunnel zone* and position and movement is continuously recorded after this time point until the end of the 30 min Exploration or Foraging Phase. The XY position of the center of the mouse is video tracked with at a rate of 30 frames per second. For the data analysis, tracking data from 0-25 minutes are used. Time spent in each zone, latency to visit a zone and number of visits to each zone, as well as the distance traveled, are calculated using the Ethovision software. The data is exported as results for the total 25-minute trial duration, as well as in 5 minute time bins. Sand displacement and food consummation measures are collected and calculated manually.

##### Behavioral Measures

All measures captured during Exploration or Foraging phases of the assay are presented in (Stacher Hörndli et al., 2019). All ‘time spent in zone’, sand displaced, food consumed and zone visit measures were normalized to the total time spent in the arena (TTA) by dividing each value by TTA (x/TTA). For time bin values, the TTA for the corresponding time bin is used for normalization (i.e., x-1/TTA-1). All latencies to visit pots were normalized to the latency to enter the platform (LEP) by subtracting the LEP (x-LEP). Latency to the center zone after arena entry (LCAE) is already normalized to LEP and does not need any further normalization. All percentage measures do not need any normalization. Time on the platform itself and time in the tunnel (cage) are not included in the normalized dataset because they are closely related and redundant to TTA.

##### Locomotor Measures

The raw data files generated by Noldus Ethovision listing all XY coordinated and all zones as well as distance traveled and velocity for each frame were used to extract data describing excursions and locomotor patterns using custom code, including the duration and number of bouts at different velocities. An excursion is defined as beginning when the mouse leaves the tunnel (In Tunnel zone) and ending when the mouse returns to the tunnel (one round trip). Continuous velocity values in the data were categorized into three velocity classes, *slow*: velocity ≤ 5 equals; *medium*: velocity > 5 ≤ 15; *fast:* velocity > 15. The length of the bout is calculated using the number of frames in the sequence and all sequences of the same velocity class longer than 3 frames are counted as a single velocity bout. In addition, a vector of several descriptive statistics is created that describes locomotor patterns and variability from X-Y tracking data.

### QUANTIFICATION AND STATISTICAL ANALYSIS

#### Foraging Behavior

##### Excursion Data Capture

Our DeepFeats approach for analyzing modularity in foraging was performed as previously described ^25^ with some modifications and advances. In our study, mice were tracked with Noldus Ethovision software. Noldus settings were used to define regions of interest in the foraging arena and indicated when the mouse was in each area. To ensure the tracking is equivalent across different mice, a Procrustes transformation of the XY coordinates was performed to put every tracking file in the same coordinate space. The track coordinates were zero’d to the center of the tunnel to the home cage. We then generated custom code in R to parse the raw Noldus tracking files into discrete, round trip home base excursions from the home cage tunnel. Each excursion is assigned a unique ID key that we call the Concise Idiosyncratic Module Alignment Report (CIMAR) string key. It stores the coordinates of the excursion in the data and the CIMAR string includes metadata regarding the mouse number, excursion number, sex, age, genotype and phase. Next, custom code compares the CIMAR coordinates to the raw Noldus data files and constructs a new dataset that extracts 57 measures from the Noldus output, which we use to initially statistically describe each excursion. The 57 measures are designed to capture a relatively comprehensive array of different behavioral and locomotor parameters, as well as describe interactions with food and non-food containing patches and exposed regions in the environment. These measures consist of shape, frequency, order and location statistics of an animal’s X and Y movements, numbers of visits and time spent at different features in the arena, including food patches (Pots#2 and 4), non-food containing patches (Pots#1 and 3), the tunnel zone, wall zone and center zone of the arena and data describing locomotor patterns, including velocity, gait and distance traveled. The 57 measures for each excursion are scaled (normalized and centered across excursions) because they are in different units.

##### Behavioral Measures to Resolve Modularity in Excursions

This section details the methods to define the set of behavioral measures that best resolve candidate modules. A data matrix was constructed in which the rows are excursions performed by the mice, labeled by CIMAR keys, and the columns are the 57 behavioral measures. Dimension reduction was performed using Principal Component Analysis and the number of principal components (PCs) to retain for the identification of modules was defined based on the set that maximized cluster identification in the training dataset partition. Unsupervised clustering was performed on the retained PCs. We used the Ward.D2 minimum variance method implemented using the “hclust” function in R to perform the clustering and define compact, spherical clusters. We then statistically define discrete excursion clusters from the results using the Dynamic Tree Cut algorithm (Langfelder et al., 2008). This is a powerful approach because it is adaptive to the shape of the dendrogram compared to typical constant height cutoff methods and offers the following advantages: (1) identification of nested clusters; (2) suitable for automation; and (3) can combine the advantages of hierarchical clustering and partitioning around medoids, giving better detection of outliers. We detect clusters using the “hybrid” method and use the DeepSplit parameter set to 4 and the minimum cluster size set to 20. The total number of clusters detected is quantified at each correlation threshold. Conceptually, more relaxed correlation threshold cutoffs could reduce cluster detection by retaining redundant measures that mask important effects from other measures. On the other hand, thresholds that are too stringent could reduce cluster detection by pruning informative measures. Our objective is to identify the threshold that uncovers the most informative and sensitive set of measures for resolving different clusters of excursions, setting the epoch for the discovery of potential modules.

##### Statistical Validation of Significant Clusters of Excursions

In our study, Dynamic Tree Cut will deeply cut branches in a dendrogram generating large numbers of small clusters if there are few bona fide relationships in the data. Thus, to test whether bona fide clusters of excursions exist in the data we implemented a random sampling procedure in R in which we randomly sample from the matrix of the retained behavioral measure data to break the relationships between the excursions and the measures. The sampled null data matrix is then subjected to the same clustering and quantification procedure to determine the number of clusters found by Dynamic Tree Cut. A null distribution is created from 10,000 iterations and compared to the observed number of clusters, which is expected to be significantly less than the null due to bona fide biological relationships between the excursions and set of retained measures. A lower tailed p value was computed to test this outcome. In a modification compared our previous study,

##### IGP Permutation Test for Stereotyped Behavioral Modules

To test whether reproducible modules of behavior exist in the data for the foraging excursions, we use the in- group proportion (IGP) statistical method for testing for reproducible clusters between two datasets ^64^. We built a modified version of this function for parallelized computing to speed the analysis for large numbers of permutations. The excursion data for the mice is separated into a training data and test data partition for reproducibility testing. A balanced partition was generated according to genotype, sex and phase factors using the “createDataPartition” function in the caret package in R. Unsupervised hierarchical clustering was performed on the Training data partition excursions and clusters were defined using Dynamic Tree Cut. Next, the centroids for each training data cluster were computed as the mean values of the behavioral data for the excursions in the cluster. The training data centroids were then used to compute the IGP statistic for each training data cluster based on the test partition data, thereby evaluating the reproducibility of each cluster.

In a modification compared to our previous study described ^25^, we created a custom IGP permutation test that is based on a distance calculation, rather than the correlation implementation in the clusterRepro R package. The distance IGP testing framework was written in C++ and speeds the permutation test by many fold and is a more accurate replication of the clustering parameters used in the test data. We used this approach to compute p values for each cluster to determine whether the IGP value is greater than chance. False positives due to multiple testing errors were controlled using the q-value method ^65^. Modules are thus defined as significantly reproducible training partition excursion clusters (q < 0.1). Each module detected was assigned an ID number and individual excursions in the data were annotated based on the module they match to. This approach facilitated quantifications of module expression frequency by the mice.

##### Statistical Modeling of Module Expression Counts

To statistically evaluate the genetic factors that significantly affect module expression frequency, we used generalized linear modeling functions implemented in R. The hypotheses tested are detailed in the text for each analysis and below. An interaction effect between module (or non-modular behavior type) was tested using the model: Expression count ∼ sex + genotype + module + genotype*module compared to a nested model of the main effects. Post-testing to define the specific modules that are affected was performed using a glm testing for a main effect of genotype on each individual module with the model: Expression count ∼ sex + genotype. We filtered based on overall variance to remove modules with low expression variance across all mice in the study and reduce multiple testing errors, which is a proven two-step method for analyzing high dimensional data ^66^. P- values were corrected for multiple testing errors by the q-value method to control the false discovery rate and pi0 was computed to determine true nulls in the data ^67^. Generalized linear modeling was performed using a Poisson distribution. We tested the goodness-of-fit of the Poisson for each module with a chi-square test of the residual deviance and degrees of freedom in R.

##### Statistical Modeling of Module Transitions

To statistically evaluate transitions between modules, the start time for each expressed module was recorded. We then constructed 1-step transition matrixes that relate the expressed module to the next expressed module to examine sequences of module expression. To determine whether module expression transitions are dependent on the identity of the previously expressed module, we performed a Fisher’s Exact test on the transition count matrix. To compare wild-type and knockout mouse transition matrices, we computed the stationary probability distribution and then the Euclidean distance between the male versus female distribution. Next, we compared the observed distance to a distribution of distances derived from randomly permuted data. In the permutation test, modules are randomly sampled from the wild-type and knockout data, two transition matrices are generated, the stationary probability distribution is computed and then compared by Euclidean distance. The observed distance is compared to the permuted distribution (>10,000 permutations) to determine the p-value.

##### Behavioral Cartography and Visualization

visualize the structure of the foraging behavior landscape and the relationships between modules and non­dular behavioral components, the raw data for 57 different behavioral measures was collected to describe h excursion for all1609 excursions performed by all mice in the study. The data were analyzed and visualized using the PHATE algorithm ^42^ and annotated according to module and non-modular components as defined using the methods *above*.

## SUPPLEMENTAL FIGURES, VIDEOS AND LEGENDS

**FIGURE S1.**
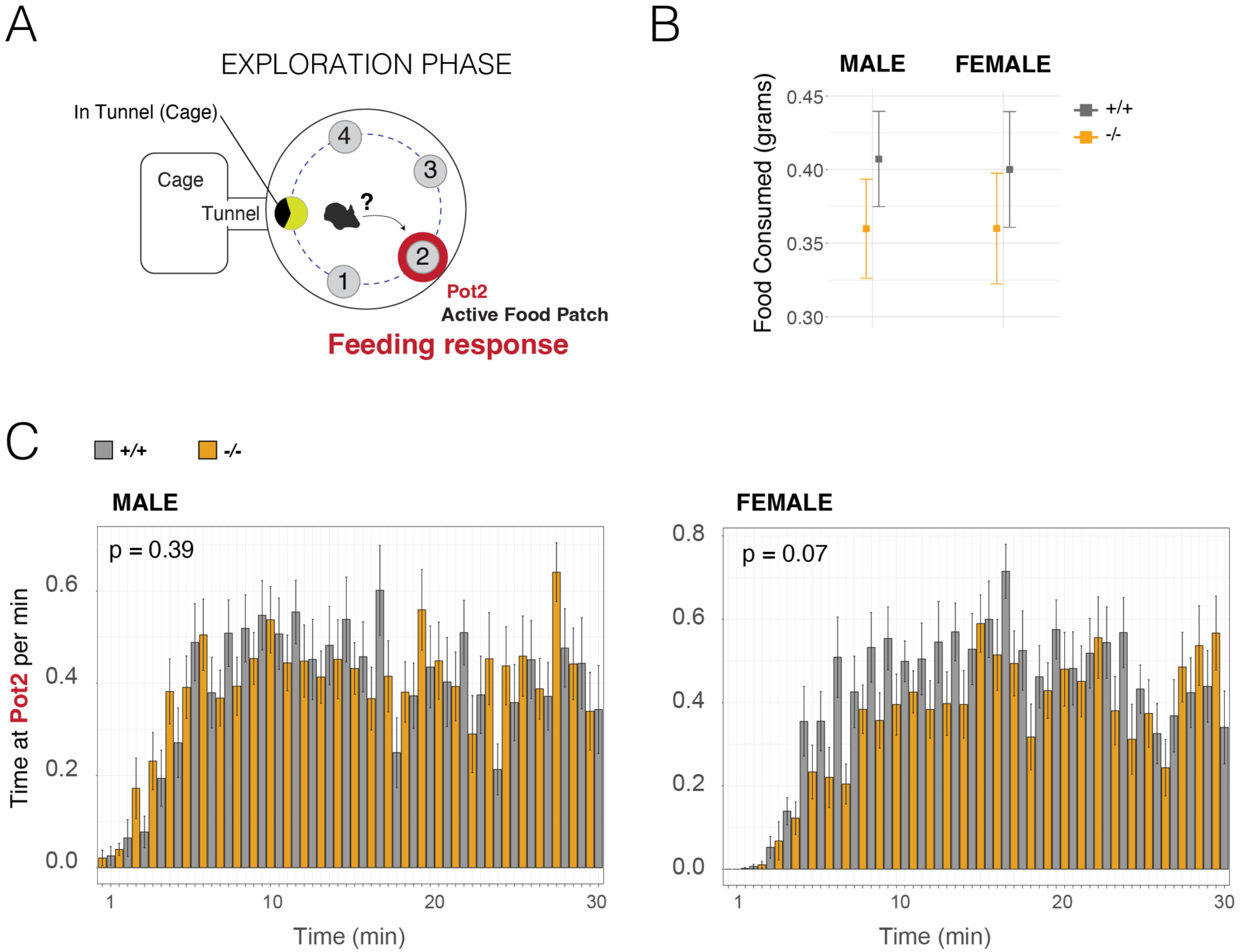
Related to Figure 2. *Arc^−/−^* mice do not show significant changes to food intake or interactions with the food patch (Pot2) during the naïve Exploration phase. (**A**) Schematic summary of the Exploration phase. (**B**) The plot shows the total food consumed by *Arc^−/−^* (orange) versus ^+/+^ (black) mice in the Exploration phase. A significant difference is not observed. N=15, One-way Anova. (**C**) The plot shows total time at the food patch during the Exploration phase (Pot2) broken down by 1 minute time bins. A significant difference is not observed between *Arc^−/−^* and *^+/+^* mice for males or females. N=15, mean+SEM, KS test comparison of distributions.

**FIGURE S2.**
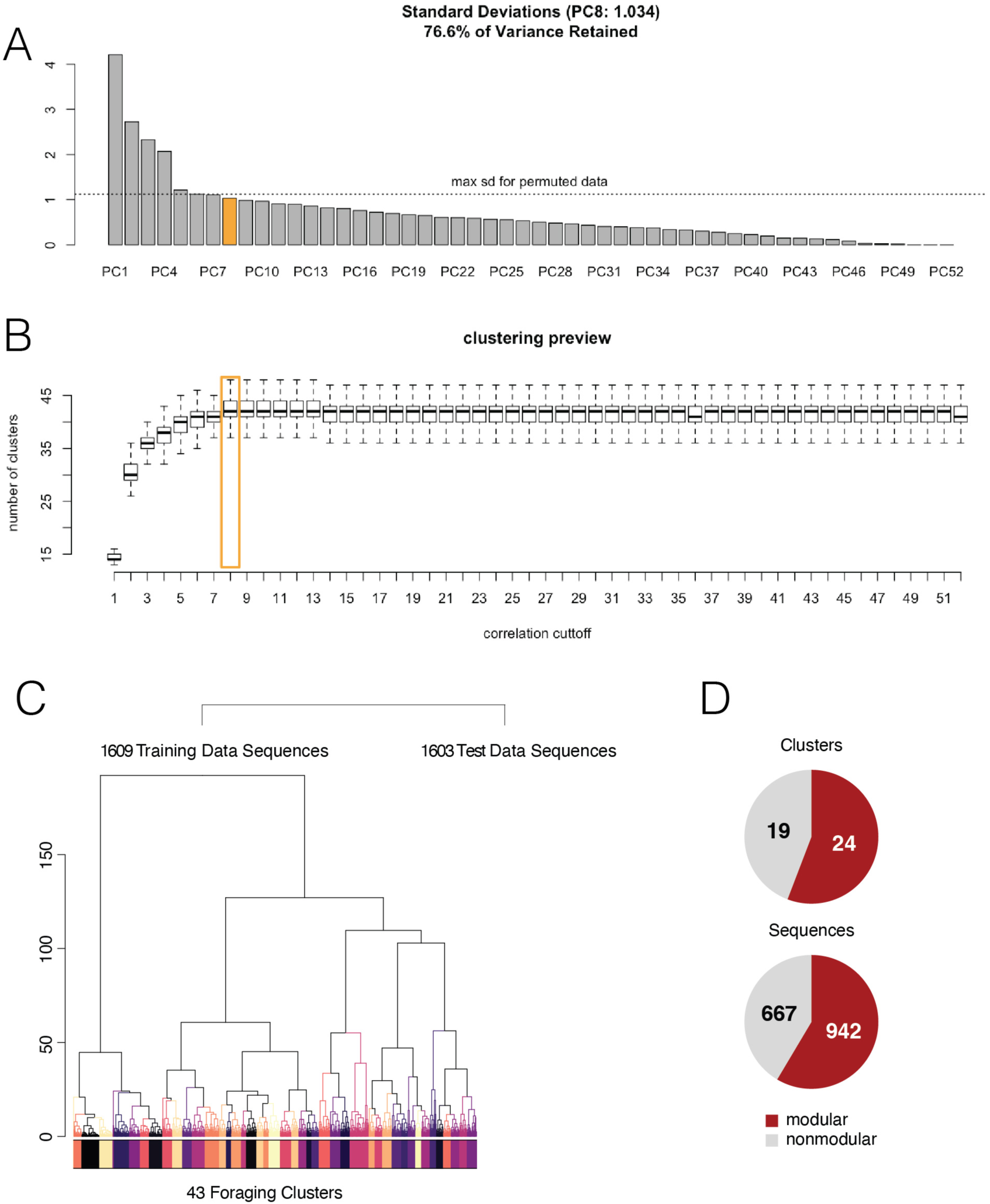
Identification of foraging modules. Related to Figure 4. (**A**) The plot shows the number of principle components (PCs) and proportion of variance explained by each PC. The chosen PC cutoff is in orange at the elbow of the plot and captures 76% of variance in the data for hundreds of measures describing discrete round trip foraging excursions from the home. (**B**) The plot shows the number of clusters discovered with the selected PCs based on different cutoffs for the number of PCs chosen. The data reveal that the maximum cluster number and minimum standard deviation in terms of cluster number is achieved using 8 PCs. (**C**) Shows the DeepFeats training and testing partitions and clustering results to identify behavioral modules. Dynamic tree cutting yielded 43 candidate clusters in the training data for module testing. (**D**) The pie charts show the number of significant modules (red) versus training set clusters that are not modular (grey) (ie. Significantly reproducible in the test data). The number of round-trip excursions found to be modular (red) versus non-modular (grey) is shown below. In-group proportion permutation test, Q-value <0.1. (**E** and **F**) The plots show representative X-Y coordinate traces of the mouse center point body movement pattern over time (Z-axis) for examples of the decision and action sequences for significant behavior modules, including two modules involving increased interactions with the food patch Pot4 (4) in the Foraging phase (E, Modules 1 and 12), compared to a module that involves increased interaction with the former food patch Pot2 (2) (F, Module 21). The traces are for the excursions in the data that are closest to the centroid of the cluster testing significant for modularity. Traces are colored by movement velocity (Black is slow or stopped and pink is moving). Mouse shadows illustrate the behavior. The dark circle is the tunnel connecting the home (H) and arena.

**FIGURE S3.**
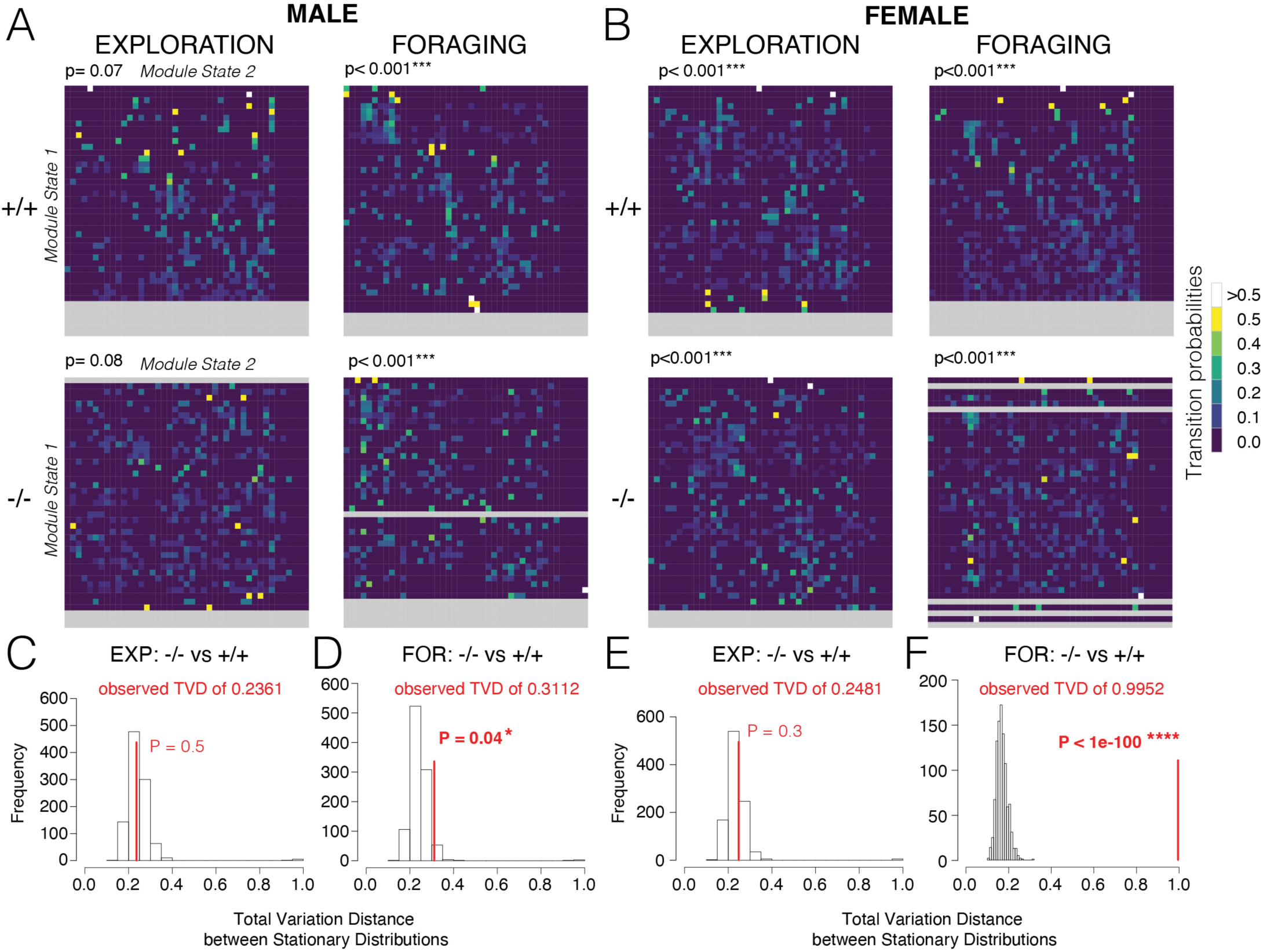
Loss of *Arc* significantly changes the sequential order of module expression in the Foraging phase. Related to Figure 4. (**A and B**) The heatmaps show transition matrices for module expression sequences during foraging behaviors in the Exploration and Foraging phase in males (A) and females (B). The y-axis shows Module State 1 and x-axis is Module State 2, where the intersections are the transition frequencies in a heatmap that shows module-to-module transition frequencies (see legend). Module expression transitions are significantly stereotyped and non-random in the Exploration and Foraging phases in *Arc^−/−^* and *+/+* mice, which is statistically shown by a Fisher’s Exact Test of independence between the rows (Module State 1) and the columns (Module State 2). The p-value shown above matrix and a significant effect indicates dependence, which is observed in all cases, though only a trend is observed in males during Exploration phase. Grey rows are modules that are not expressed due to being specific to either the Exploration or Foraging phases, or not expressed by a particular genotype. (n=15) (**C-F**) The plots show results of a permutation test comparing the transition matrices between Arc +/+ versus −/− mice in the Exploration (EXP, C and E) and Foraging (FOR, D and F) phases for males (C and D) and females (D and F). The permutation test compares the stationary probability distributions of the transition matrices for *Arc^−/−^* and *^+/+^* mice. The results show a significant difference in module expression transitions between *Arc^−/−^* and *^+/+^* mice specifically in the Foraging phase in males and females. Thus, *Arc* affects the sequential order of module expression in ^−/−^ versus ^+/+^ mice. TVD, total variation distance between transition matrix stationary distributions.

**FIGURE S4.**
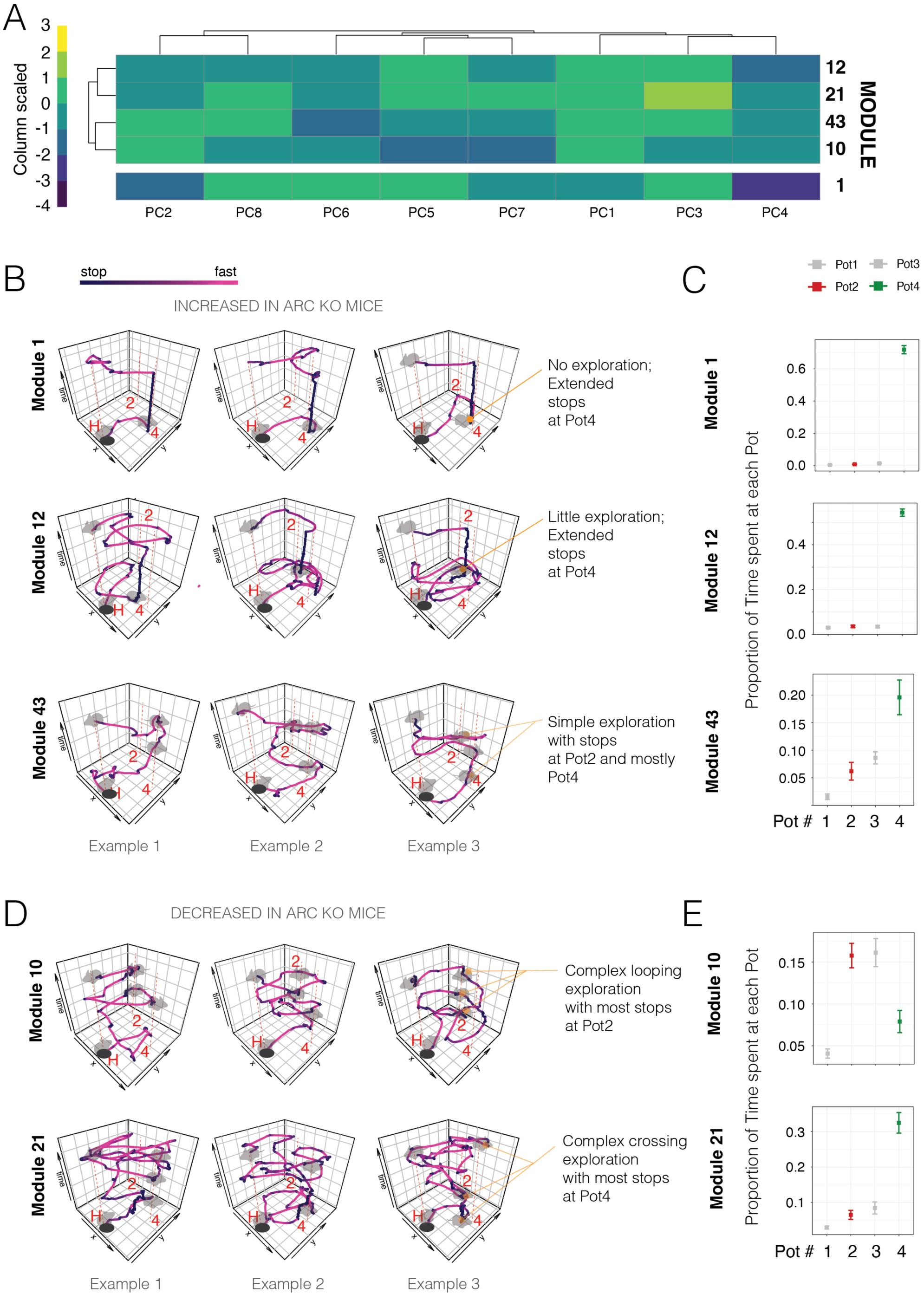
Characterization of Arc affected foraging modules. Related to Figure 4. (**A**) The heatmap shows the coordinate values for each principal component from a PCA analysis of the behavior data for the centroid behavior that is representative of each shown module. The result shows how the different PCs differentiate different modules. (**B and C**) The X-Y traces over time in (B) show three representative movement patterns from different mice for Modules 1, 12 and 43, which are modules that show significantly increased expression by *Arc^−/−^* mice. The locations of the home (H), former food patch Pot2 (2) and active food rich patch Pot4(4) are shown. Mouse stopping positions are illustrated by grey mice. Module 1 and 12 show simple decision and action sequences that are focused on the food containing patch in Pot4. Module 43 involves a simple exploration with brief stops at Pot4 and Pot2. The graphs in (C) show the average proportion of time spent at each pot for each module type across different mice. Modules 1 (n= 43 mice), Module 12 (n=44 mice) and Module 43 (n=19) show that most time is spent at Pot4. Module 43 also involves time at the former food patch, Pot2. Plots show mean ± SEM. (**D** and **E**) The X-Y traces show representative movement patters for three different mice for Modules 10 and 21 (D), which have significantly decreased expression in *Arc ^−/−^* mice compared to ^+/+^ littermates. Both modules are complex compound decision and action sequences that involve explorations and interactions with different elements of the environment. Module 10 has a simpler looping structure compared to Module 21. The graphs in (E) show the proportion of time spent at each pot, revealing that Module 10 (n=26) involves increased time at the former food patch (Pot2), while Module 21 involves time at Pot4 with some time at Pot2 (n=34).

## SUPPLEMENTAL VIDEOS

**Supplemental Video S1. 3D PHATE projection of the multi-dimensional data for 1609 foraging excursions performed by male and female *Arc^−/−^* and ^+/+^ mice.** Each dot represents the data for one excursion. Colors show the excursions for different modules. Grey dots are non-modular excursions. Arms and branch connections in the map show the structure, relationships and similarity connections between the different excursion behaviors expressed by mice.

<u>Click to download and watch:</u>

https://.dropbox.com/s/08r3h45zbs911jr/SupplementaryVideo_S1_ModularVersusNonmodularBehavior.mp4?dl=0

**Supplemental Video S2. 3D PHATE projection of the multi-dimensional data for the excursion data for modules only shows clear separation and relative relationships within the structure of the foraging behavior landscape for mice.** Each dot represents the data for one excursion.

Different colors show the excursions for different modules. Arms and branch connections in the map show the structure, relationships and similarity connections between the different modules expressed by mice.

<u>Click to download and watch:</u>

https://.dropbox.com/s/92ro9w3ht5fqu08/SupplementaryVideo_S2_ModularOnly.mp4?dl=0

**Supplemental Video S3. 3D PHATE projection of the multi-dimensional data for 1609 foraging excursions performed by male and female *Arc^−/−^* and ^+/+^ mice showing the subset of modules that are significantly affected by loss of *Arc*.** Each dot represents the data for one excursion.

Different colors show the different *Arc* affected modules. Grey indicates excursions and behaviors that are not significantly affected in *Arc^−/−^* mice. Arms and branch connections in the map show the structure, relationships and similarity connections between the different modules affected by loss of *Arc*.

<u>Click to download and watch:</u>

https://.dropbox.com/s/2mz5aujrkl2urj8/SupplementaryVideo_S3_Arc_affected_modules_red%20is%201%2C%20cyan%20is%202%2C%20purple%20is%2010%2C%20blue%20is%2012%2C%20yellow%20is%2021%2C%20green%20is%2043.mp4?dl=0

